# STATegra: a comprehensive multi-omics dataset of B-cell differentiation in mouse

**DOI:** 10.1101/587477

**Authors:** David Gomez-Cabrero, Sonia Tarazona, Isabel Ferreirós-Vidal, Ricardo N. Ramirez, Carlos Company, Andreas Schmidt, Theo Reijmers, Veronica von Saint Paul, Francesco Marabita, Javier Rodríguez-Ubreva, Antonio Garcia-Gomez, Thomas Carroll, Lee Cooper, Ziwei Liang, Gopuraja Dharmalingam, Frans van der Kloet, Amy C. Harms, Leandro Balzano-Nogueira, Vicenzo Lagani, Ioannis Tsamardinos, Michael Lappe, Dieter Maier, Johan A. Westerhuis, Thomas Hankemeier, Axel Imhof, Esteban Ballestar, Ali Mortazavi, Matthias Merkenschlager, Jesper Tegner, Ana Conesa

## Abstract

Multi-omics approaches use a diversity of high-throughput technologies to profile the different molecular layers of living cells. Ideally, the integration of this information should result in comprehensive systems models of cellular physiology and regulation. However, most multi-omics projects still include a limited number of molecular assays and there have been very few multi-omic studies that evaluate dynamic processes such as cellular growth, development and adaptation. Hence, we lack formal analysis methods and comprehensive multi-omics datasets that can be leveraged to develop true multi-layered models for dynamic cellular systems. Here we present the STATegra multi-omics dataset that combines measurements from up to 10 different omics technologies applied to the same biological system, namely the well-studied mouse pre-B-cell differentiation. STATegra includes high-throughput measurements of chromatin structure, gene expression, proteomics and metabolomics, and it is complemented with single-cell data. To our knowledge, the STATegra collection is the most diverse multi-omics dataset describing a dynamic biological system.

## Background and Summary

The concept of multi-omics and data-integration has been increasingly used during the last 5 years to describe the multitude of high-throughput molecular technologies that can be applied to the study and analysis of biological systems^1^. Such techniques hold the promise to uncover the different biological processes and layers of regulatory complexity within biological systems. In brief, high-throughput molecular methods can extract information of essentially three basic, yet different components of living cells. Nucleic acids can readily be profiled using massive, parallel sequencing, which in turn provide deep a characterization of chromatin properties (i.e. Hi-^2^C, ATAC-seq^3^, DNase-seq^4^, ChIP-seq^5^, WGBS^6^, RRBS^7^) and the dynamics of gene expression (i.e. RNA-seq^8^, microRNA-seq^9,10^, PAR-CLiP^11^, iCLIP-seq^12^). Proteins are measured by proteomics and phosphoproteomics approaches, based on Liquid Chromatography (LC) or Isotope-coded affinity tag labeling (iTRAQ) coupled to Mass Spectrometry (MS). Finally, the metabolome and lipidome, i.e. organic compounds, are captured using mature techniques such as LC/GC-MS or Nuclear Magnetic Resonance (NMR). Increasingly, multi-omics technologies are applied during the same physiological conditions from either the same or different samples to generate a comprehensive set of data spanning multiple molecular levels. The general expectation of multi-omics projects is that the combination of multi-layered data will reveal aspects of the complexity of biological systems that cannot be fully understood using only a particular data-type. Moreover, in addition to the exciting technical reality of being able to monitor several complementary data-types, the community has come to realize the power of using time in the experimental design. Hence, by collecting data over time, where as a rule the different molecular entities are correlated, it is much more amenable to extract key processes from each data-type as well as uncovering dependencies between different regulatory layers. These technical and conceptual advances are currently being transferred into the vibrant single-cell biology community. Thus, recent advances in single-cell omics technologies have made it feasible to perform multi-omics profiling of individual cells. Consequently, the single-cell community can benefit from the experiences and lessons derived from time-dependent bulk multi-omics analysis. Clearly, a high-resolution single-cell analysis has proven crucial to assess tissue heterogeneity^13–15^, cell fate^16,17^. In conclusion, we are most likely entering an era where we can target regulatory networks in single cells^18^ using a temporal paradigm coupled to a multiomics analysis.

While multi-omics projects are frequently depicted as a set of stacked molecular layers that are connected to pass information from the genetic component to the organismal phenotype, the harsh reality is that still many multi-omics project are constrained by budgetary restrictions and sample limitations which evidently reduce the number technologies that can realistically be assessed. In most cases, only a few data types can be included, with a limited number of samples, and analyses is as a rule restricted to focus on 2 or 3 regulatory layers. A few international projects have however successfully collected large datasets and generated comprehensive portfolios of omics measurements. For example, ENCODE^19^, TCGA^20^, IHE^21^, ImmGen^22^, had the explicit goal to perform an extensive characterization of a particular set of cells or tissues. These projects have impacted the scope and type of analysis methods and scientific discoveries that can be achieved so far by the multi-omic approach. In some cases combining multi-level data has the ambition to increase the required statistical power to enable the classification of samples or predict disease outcomes. By measuring different types of features the chance of identifying relevant biomarkers increases, but the analysis does not automatically lend itself to a mechanistic account of the inter-dependencies between these biomarkers as well as their relationship with the outcome, such as a disease. In some cases however, two specific omics layers are measured in order to probe their regulatory relationships. For example, methods that integrate ATAC-seq or RRBS with RNA-seq might shed light on the epigenetic control of gene expression^23^, while integrating transcriptomics and metabolomics data may help elucidate metabolic regulation^24,25^. Yet, there have been very few multi-omic studies that evaluate dynamic processes such as cellular growth, development and adaptation. Hence, we still lack formal analysis methods and comprehensive multi-omics datasets that can be leveraged to develop true multi-layered models for dynamic cellular systems. This state-of-affairs has been the rationale underpinning the formulation of what is referred to as the STATegra project (http://www.stategra.eu/). This is a transnational initiative to develop methods, software and data for dynamic multi-omics analyses. From the STATegra project several tools for integrative multi-omics data analyses have been published and released^26–33^.

Here we share the collection of the different STATegra datasets, a multi-omics dataset that combines measurements from up to 10 different omics technologies applied to the same biological system. STATegra uses a well-studied system, namely mouse pre-B-cell differentiation, in a cell line model^34^. This is a highly reproducible *in vitro* system^33–36^ that allows the generation of sufficient material to deploy a comprehensive set of omics measurements. STATegra covers the three types of biomolecules and the different layers that comprise the basic flow of genetic information: chromatin structure (through DNase-seq, RRBS and ChIP-seq), gene expression (RNA-seq and miRNA-seq), proteomics and metabolomics. The collection is complemented with single-cell RNA-seq and ATAC-seq data on the differentiating conditions. The STATegra multi-omics dataset is unique in the number and diversity of omics technologies available and in the dynamic nature of the system. Our ambition has been to generate this collection of data to serve – in full or using parts of it-as workbench for the development of integrative analysis methods for the multi-layered systems biology.

In previous studies, ChIP-seq data from this collection have been used to identify Ikaros targets^34^. ChIP-seq, DNase-seq, RNA-seq and scRNA-seq datasets were used in Vidal *et al.*^37^ to describe the cross-talk between IKAROS Foxo1 and Myc transcription factors in regulating B-cell development. scATAC-seq, scRNA-seq and ATAC-seq data have been used to develop new statistical methods for the integration of single-cell multi-omics^33^.

## Methods

### Experimental design

Figure 1 illustrates the STATegra dataset. The mouse B3 cell line models the pre-BI (or Hardy fraction C’) stage. Upon nuclear translocation of the Ikaros transcription factor these cells progress to the pre-BII (or Hardy fraction D) stage, where B cell progenitors undergo growth arrest and differentiation^34,38^. The B3 cell line was retrovirally transduced with a vector encoding an Ikaros-REt2 fusion protein, which allows control of nuclear levels of Ikaros upon exposure to the drug Tamoxifen^34^. In parallel, cells were transfected with an empty vector to serve as control for the Tamoxifen effect. After drug treatment, cultures were harvested at 0h, 2h, 6h, 12h, 18h and 24hs (Figure 1A) and profiled by several omics technologies: long messenger RNA-seq (mRNA-seq) and micro RNA-seq (miRNA-seq) to measure gene expression; reduced representation by bisulfite sequencing (RRBS) to measure DNA methylation; DNase-seq to measure chromatin accessibility as DNaseI Hypersensitive Sites (DHS) and transcription factor footprints, shotgun proteomics and targeted metabolomics of primary carbon and amino-acid metabolism. Moreover, single-cell RNA-seq (scRNA-seq) data for the entire time-series, while bulk ATAC-seq (ATAC-seq) and single-cell ATAC-seq (scATAC-seq) were obtained in a later round of experiments for 0h and 24h-time points of Ikaros induction only (no control series were run for these datasets). The dataset is complemented by existing ChIP-seq data on the same system equivalent to our 0h and 24h time points^34^. In total, 793 different samples across the different omics datasets define the STATegra data collection (Figure 1B).

**Figure 1.**
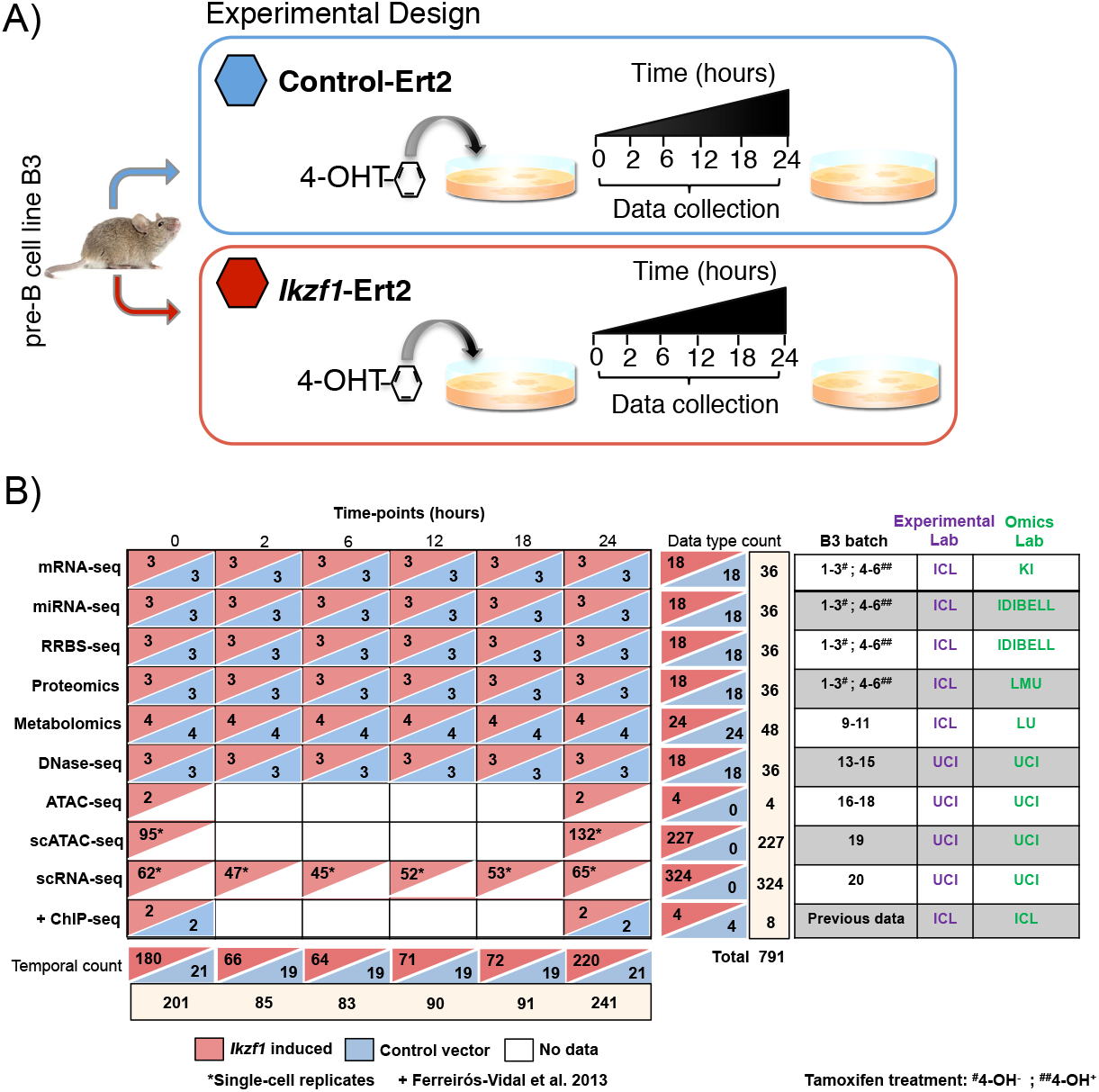
STATegra data generation. A) Inducible Ikaros B3 cell system. Time course experiment collects samples at 6 time-points after Tamoxifen induction of Ikaros expression, Control cells carry empty vector. B) Diversity of omics platforms, number of biological replicates, batch distribution and lab assignment for B3 cell culture and omic library preparation. Data on each row corresponds to the one omics type on the left. + Previous data from ^34^.

The time points analyzed were based on previous microarray studies^34^ and have been fully validated by comparing the transcriptional response in this experimental system to pre-B cell differentiation in vivo^37^. Ikaros translocates to the nucleus of B3 cells within minutes, binds to target promoters and changes RNAP2 occupancy and primary transcript levels with immediate effect^36^. The 2h time point is relatively late compared to changes in primary transcript levels^36^ and was chosen because the data presented here were generated by conventional RNA-seq, which relies on changes in steady state, rather than primary transcript levels.

### Culture conditions

B3 cells containing inducible Ikaros can be expanded before induction of Ikaros to produce sufficient material for all omics experiments. G1 arrest occurs within 16 h following Ikaros induction. Cells containing inducible Ikaros were generated by transducing mouse pre-B cell line B3 with mouse stem cell virus (MSCV) retroviral vectors encoding a fusion protein of haemagglutinin-tagged wild type Ikaros (HA-Ikaros) and the estrogen receptor hormonebinding domain (ERt2), followed by an internal ribosomal entry site (IRES) and GFP. Control cells were generated by transducing mouse pre-B cell line B3 with mouse stem cell virus (MSCV) retroviral vectors encoding the estrogen receptor hormone-binding domain (ERt2) followed by an internal ribosomal entry site (IRES) and GFP. Retroviral infected B3 cells were sorted based on GFP levels. GFP positive cells were expanded in culture for few days (3-4) and then frozen. Frozen vials containing 5 million cells were stored in liquid nitrogen.

For time course experiments, 10 million control and Ikaros cells were thawed and expanded for 4 days. Four days later cells were plated for induction of the different time points. Both control and Ikaros cells were split in flasks containing 20 million cells at a density of 0.5 million cells per ml each. For time point inductions, 0.5uM 4-hydroxy-tamoxifen (4-OHT) was added to both, a flask containing Ikaros cells and a flask containing control cells, at one of the specified times: 2h, 6h, 12h, 18h or 24h before collection. Cells for time point 0h (no 4-OHT) induction were obtained separately in three different batches (Figure 1B). All cells within the same experimental batch were harvested simultaneously. Cells were centrifuged for 5 min at 1200 rpm, washed twice in PBS and counted to aliquot. Aliquots of 10 million cells were done for RNA-seq and metabolomics and proteomics platforms and of 5 million cells for miRNA-seq and Methyl-seq platforms. Cell pellets were snap-frozen in liquid nitrogen and stored at −80. 20-25 million and 50,000 cells were used for DNase-seq and bulk ATAC-seq samples. The full time course experiment was repeated different times (batches) to generate biological replicates (Figure 1B). The same physical cultures were used to obtain cells for mRNA-seq, miRNA-seq, RRBS and proteomics. Other omics technologies ran their own cultures to obtain cell material.

### Acquisition of Multi-omics data

#### RNA-seq

Total RNA was isolated with RNAbee (Ambion), frozen ICL and transported *via courier* (<1 day) to Karolinska Institutet. To account for the impact of the different sources of variability during RNA-seq profiling, we implemented a carefully balanced distribution of samples in relation to time points (6 time points), treatment (Ikaros vs Control), library preparation, barcode, sequencing run and lanes and biological replicates (3 batches). Briefly, samples were first balanced in six library preparation runs of 6 samples each (Figure 2). Secondly, each RNA-seq library was split into two (total of 72) in order to better account for variability associated with sequencing. Finally, for sequencing, 75 nucleotides paired-end, the 72 libraries were balanced into 4 flow-cells and in each lane we included 3 libraries. In each lane, we ensured to have different libraries, different batches, different time points and at least both conditions present. Additionally, we balanced the time-points, conditions and batches within each flow-cell. For each flow-cell, a full lane was reserved for quality control. We aimed to obtain 50M reads per library, therefore 100M reads per sample. Libraries were built using the strand-specific RNA-seq dUTP protocol^39^. Sequencing was conducted on an Illumina HiSeq 2500 platform.

**Figure 2.**
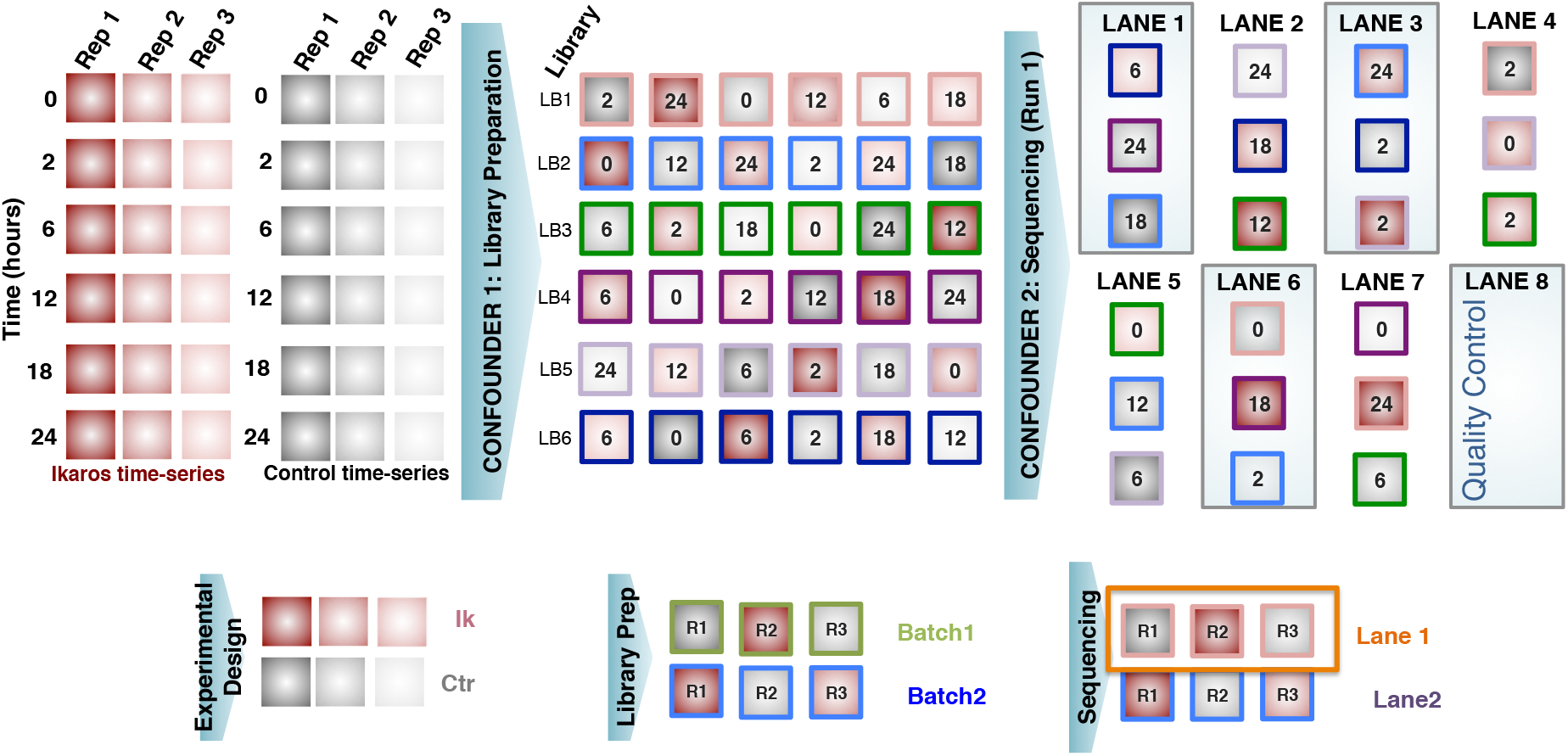
Experimental design for RNA-seq.

#### Small RNA-seq for miRNA analysis

Small RNA-seq analysis was performed using Trizol-extracted total RNA of 3 biological replicates (4,5,6) for time 0h and total RNA of 3 biological batches (1, 2 and 3) for times 2h, 6h, 12h, 18h and 24h. RNA quality was assessed using Bioanalyzer (Agilent Technologies) evaluating the RNA integrity number (RIN). The library was generated using TruSeq Small RNA Sample Preparation Kit and deep sequencing was performed in Illumina Hiseq 2000 platform. Between 15 and 20 millions of sequencing reads were obtained from each sample.

The library preparation and sequencing of the biological replicates were conducted in two different occasions (technical batches). Figure 3 shows the experimental design according to the batch in which samples were processed. There were two experimental conditions (C=Control, IK=Ikaros) and the 3 biological replicates per condition and time point were numbered as 1, 2 and 3. For some of these biological replicates one additional technical replicate was generated (Figure 3) in order to estimate the variability between technical batches and to correct any potential batch effect.

**Figure 3.**
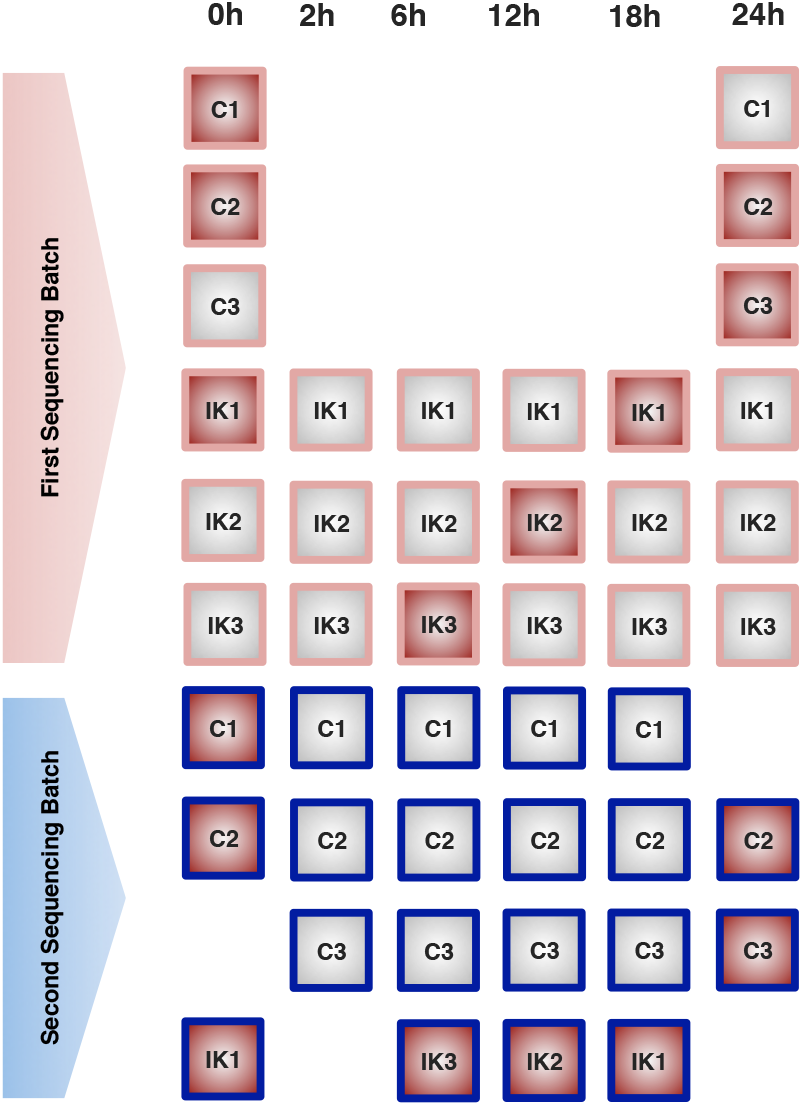
Experimental design for small RNA-seq. Two sequencing batches were run. Samples with red filling were repeated at both batches to allow for estimation of batch effects.

#### DNase-seq

DNase-seq was performed on ~20-25 million cells with 3 biological replicates for all time-points (0-24 hours) and conditions (Ikaros-inducible and control). Briefly, cells were harvested and washed with cold 1X PBS, prior to nuclei lysis. Lysing conditions were optimized to ensure >90% recovery of intact nuclei. DNaseI concentrations were titrated on Ikaros-inducible and control cells using qPCR against known positive DNaseI hypersensitive promoters (Ap2a1, Ikzf1, Igll1) and negative inaccessible hypersensitive promoters (Myog, Myod) in our biological system, thereby reducing excessive digestion of DNA. Enrichment of DNaseI hypersensitive fragments (0-500bp) was performed using a low-melt gel size selection protocol. Library preparation was performed and sequenced as 43bp paired-end NextSeq 500 Illumina reads. DNaseI libraries were sequenced at a minimum depth of 20 million reads per each biological replicate. To perform DNaseI footprinting analysis, libraries were further sequenced and merged to achieve a minimum of 200 million mapped reads.

#### RRBS

Genomic DNA was isolated using the high salt method and used for reduced representation bisulfite sequencing (RRBS), a bisulfite-based protocol that enriches CG-rich parts of the genome, thereby reducing the amount of sequencing required while capturing the majority of promoters and other relevant genomic regions. This approach provides both single-nucleotide resolution and quantitative DNA methylation measurements. In brief, genomic DNA is digested using the methylation-insensitive restriction enzyme MspI in order to generate short fragments that contain CpG dinucleotides at the ends. After end-repair, A-tailing and ligation to methylated Illumina adapters, the CpG-rich DNA fragments (40–220 bp) are size selected, subjected to bisulfite conversion, PCR amplified and then sequenced on an Illumina HiSeq2500 PE 2×100bp^40^. The libraries were prepared for 100-bp paired-end sequencing. Around 30 million sequencing reads were obtained from each sample.

#### Single-cell RNA-seq

Single cells were isolated using the Fluidigm C1 System. Single-cell C1 runs were completed using the smallest IFC (5-10 um) based on the estimated size of B3 cells. Briefly, cells were collected for each time-point at a concentration of 400 cells/µl in a total of 50 µl. To optimize cell capture rates on the C1, buoyancy estimates were optimized prior to each run. Our C1 single-cell capture efficiency was ~75-90% across 8 C1 runs. Each individual C1 capture site was visually inspected to ensure single-cell capture and cell viability. After visualization, the IFC was loaded with Clontech SMARTer kit lysis, RT, and PCR amplification reagents. After harvesting, cDNA was normalized across all libraries from 0.1-0.3 ng/µl and libraries were constructed using Illumina’s Nextera XT library prep kit per Fluidigm’s protocol. Constructed libraries were multiplexed and purified using AMPure beads. The final multiplexed single-cell library was analyzed on an Agilent 2100 Bioanalyzer for fragment distribution and quantified using Kapa Biosystem’s universal library quantification kit. The library was normalized to 2 nM and sequenced as 75bp paired-end dual-indexed reads using Illumina’s NextSeq 500 system at a depth of ~1.0-2.0 million reads per library. Each Ikaros time-point was performed once, with the exception of 18 and 24 hour time-points, in which two C1 runs were required in order to achieve approximately ~50 single-cells per each time-point.

#### Bulk and single-cell ATAC-seq

Single-cell ATAC-seq was performed using the Fluidigm C1 system as done previously^41^. Briefly, cells were collected for 0 and 24-hours post-treatment with tamoxifen, at a concentration of 500 cells/µl in a total of 30-50 µl. Additionally, 3 biological replicates of ~50,000 cells were collected for each measured time-point to generate bulk ATAC-seq measurements. Bulk ATAC-seq was performed as previously described^3^. ATAC-seq peak calling was performed using bulk ATAC-seq samples. ATAC-seq peaks were then used to estimate the single-cell ATAC-seq signal. Our C1 single-cell capture efficiency was ~70-80% for our pre-B system. Each individual C1 capture site was visually inspected to ensure single-cell capture. In brief, amplified transposed DNA was collected from all captured single-cells and dual-indexing library preparation was performed. After PCR amplification of single-cell libraries, all subsequent libraries were pooled and purified using a single MinElute PCR purification (Qiagen). The pooled library was run on a Bioanalyzer and normalized using Kappa library quantification kit prior to sequencing. A single pooled library was sequenced as 40bp paired-end dual-indexed reads using the high-output (75 cycle) kit on the NextSeq 500 from Illumina. Two C1 runs were performed for 0 and 24-hour single-cell ATAC-seq experiments.

#### Proteomics

A heavy-isotope labeled cell line representing the preB3 cell line at the starting condition was spiked to the sample before trypsin digestion to balance differences in sample amount resulting from sample preparation. After tryptic digestion, proteomic measurements of the 36 biological batches were analyzed by one-dimensional nanoRP-C18 LC-MS/MS in technical triplicates on an LTQ Orbitrap platform coupled to an Ultimate 300 RSLC system (Thermo-Fisher). First, peptide mixtures were desalted on a trapping column (0.3×5mm, Acclaim PepMap C18, 5 µm, Thermo-Fisher) at a flow rate of 25 µl/min of 0.05% TFA. A linear gradient from 3% B to 32% acetonitrile in 0.1% formic acid in 4 h was applied optimal separation of the complete proteome sample. Peptides eluting from the column were directly transferred to the gas phase via a nano-electrospray ionization source (Proxeon) and detected in the mass spectrometer. A data-dependent acquisition cycle consisting of 1 survey scan at a resolution of 60,000 and up to 7 MS/MS scans were employed. Orbitrap MS spectra were internally calibrated on the siloxane signal at 442.1 m/z Charge-state detection was enabled allowing for a precursor selection of charges 2-5 and excluding precursors with undefined, single and higher charge. Precursors with minimal signal intensity of 5000 cps, were isolated within a 1.2 Da window and fragmented by CID (normalized collision energy 35, activation time 30 ms, Q 0.25) and analyzed in the ion trap. Previously analyzed precursors were dynamically excluded from MS/MS selection for 180 seconds.

#### Metabolomics

Metabolomics measurements were performed on different biological batches than the other omics platforms because the sample preparation part for metabolomics is different than for the rest. In particular, metabolomics requires acute stopping of all metabolic reactions after sampling, while for other types of measurements this is not so critical. The cell extraction protocol for metabolomics consisted of filtration, washing, and quenching steps to remove medium from the cells and stop metabolism. Four biological batches (9, 10, 11 and 12) were acquired. Visual inspection of the cell pellets showed that batch 11 and 12 contained samples that were not completely dry. The metabolomics measurements were obtained with two different analytical platforms, a targeted liquid chromatography mass spectrometry (LC-MS) platform and gas chromatography mass spectrometry (GC-MS) platform. The LC-MS is a targeted platform measuring amino acids and biogenic amines and the GC-MS focuses on polar metabolites of the primary metabolism such as glycolysis, cyclic acid cycle and amino acid metabolism. LC-MS and GC-MS data had measurements for respectively 36 and 40 metabolites. The measurements were done on exactly the same samples. 80% of the pooled extract was for GC-MS, 10% for LC-MS, 8% for protein weight. Some metabolites were measured at both platforms. In that case, the LC-MS value was selected.

The metabolomics measurement pipeline includes two types of control: the quality control (QC) sample and the internal standard solution. The QC sample is typically a mixture of study samples that is inserted after each six study samples in the measurement series and is used to correct for experimental drift of the analytical instrument. Because of the limited availability of sample material the QC sample used here was not a mixture of study samples but material of control B3 cells not activated with tamoxifen. The internal standard solution for the GC-MS and LC-MS consists of 13C labeled yeast extract, which is added to each study sample at the beginning of the sample preparation process to correct for experimental errors made during the sample processing. For LC-MS an additional internal standard solution is added consisting of 13C labeled amines for most of the amines measured with the platform. For LC-MS the labeled versions of the metabolites were used as internal standard while for GC-MS the best internal standard was chosen based on the smallest residual standard deviation of the QC samples. During the process of measurement, the time points for each batch were randomized, but each Ikaros sample and its control were maintained together.

### Omics pre-processing

Data pre-processing is next described in detail for each omics type. Figure 4 shows a comparative overview of the different preprocessing pipelines.

**Figure 4.**
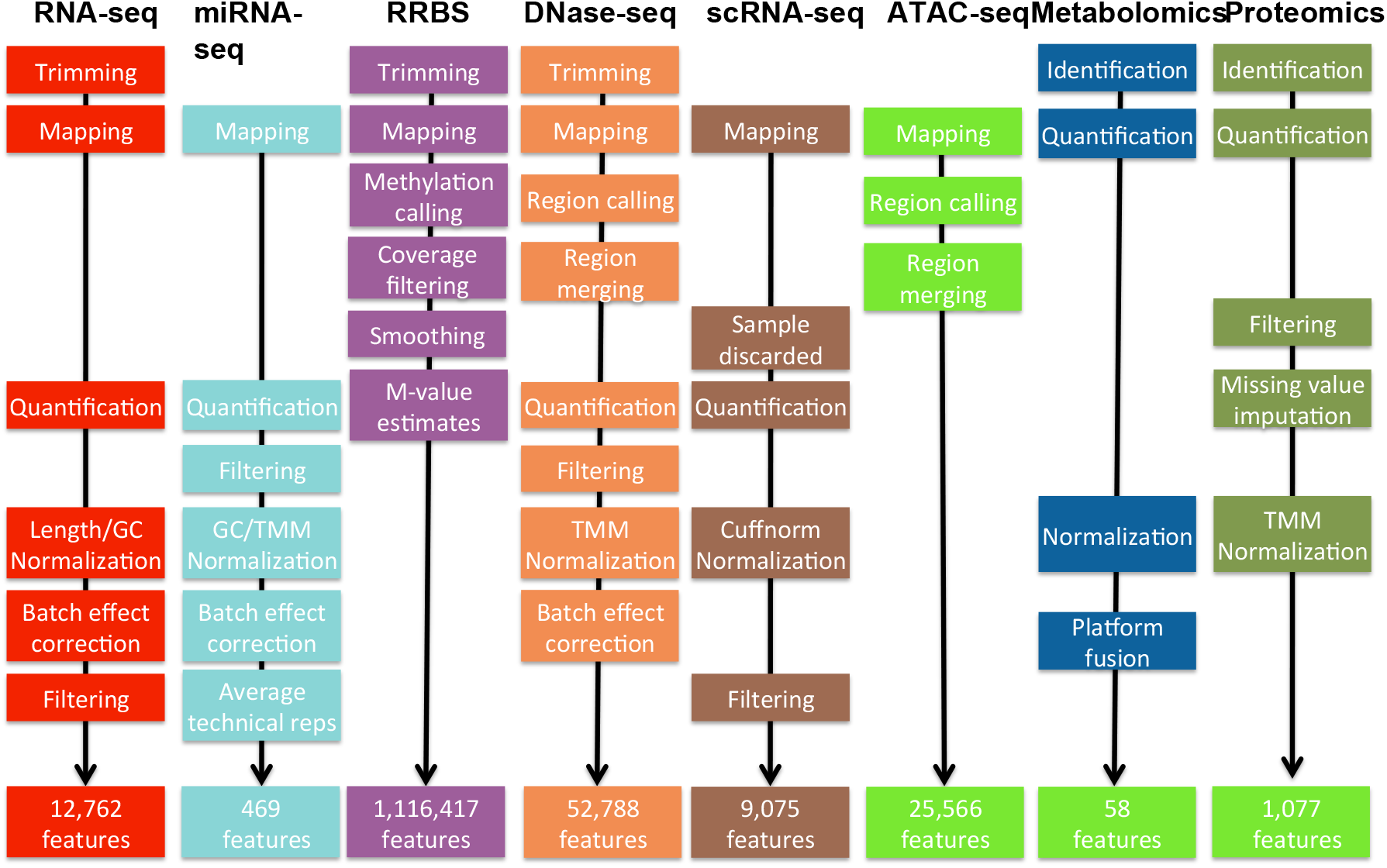
Preprocessing pipelines for 7 omics technologies. See methods for details.

#### RNA-seq

Tophat2^42^ was used to map fragments to the mm10 reference genome; the very-sensitive mode only allowing a unique best mapping per fragment was used. Piccard (https://broadinstitute.github.io/picard/) and Fastqc^43^ were used to perform a quality control considering elements such as duplication levels, GC content and k-mer overrepresentation. We observed that the duplication level was high (over 90%) in most samples as expected for high sequencing-depth in RNA-seq; additionally, some samples were having a GC content over-representation (Supplementary file S1). Trimming was applied to remove Illumina primers and low-quality nucleotides. HTSEQ^*44*^ i*ntersection-option* was used to assign fragments to genes. Data were normalized using *cqn^45^*, which corrects for GC content and gene-length. A non-parametric version of Combat methodology^46^ was used after cqn to correct for library-preparation effects.

#### miRNA-seq

The quality of the sequencing reads was checked with the Fastqc tool with good results (Supplementary file S1). Alignment of raw data was performed using Novoalign (http://novocraft.com/) on mouse miRNA sequences from mirBase. Quantification was performed using multiBamCov^47^ and counts were found for 1,086 out of 1,908 miRNAs present in the database. Low count miRNAS were further filtered out with the CPM (counts per million) method in NOISeq R package^48^ by setting a threshold of 1 for CPM. The final dataset contained 469 miRNAs. GC content bias was eliminated with the cqn R package^45^, and data was normalized by TMM^49^. PCA analysis indicated that, although Control and Ikaros samples separated well above batches, a batch effect was observed for different Ikaros time points (not shown). This bias was corrected by ComBat^46^ and technical replicates were used to avoid confounding batches with experimental conditions. After batch correction, technical replicates were averaged for further analyses.

#### DNase-seq

DNase-seq reads were trimmed to 36bp and paired-end mapped to the mm10 reference genome using Bowtie2^50^ with options: -v 2 -k 1 -m 1 --best –strata. DNase-seq peaks were called for each replicate using the HOMER findPeaks function. We employed a specific peak-calling strategy to capture several features of our DNaseI hypersensitive sites (DHS). Our strategy was to include both ‘narrow’ and ‘broad’ DHS peaks in our analysis. This captured a comprehensive set of sites with a wide DHS dynamic range. Initially, we used HOMER to determine narrow DHS peaks using a default size parameter (120-150bp) with a minimum peak distance of 50bp between DHS and an FDR of 1%. We then included a second round of peak calling, restricting to a peak size of 500bp with a minimum peak distance of 50bp between DHS peaks and an FDR of 1%. We then merged the two peak sets for each replicate. We required a minimum 1bp overlap of peaks across all three biological replicates for each time-point respectively and generated a consensus DHS peak list across all time-points.

The consensus DHS (53,624) were filtered for chrM peaks, partial chromosomes, and mouse ENCODE blacklist regions. Counts representing the chromatin accessibility were estimated for each consensus DHS using the Bedtools coverageBed function. Additionally, no DHS were considered with less than 10 reads (~1 RPM) in all time-points, resulting in a final dataset with 52,788 consensus DHSs. Data were normalized by a combination of RPKM and TMM. An unwanted source of variability was detected in the data that could not be associated with any experimental factor such as production batches or library preparation. Therefore, a method like ComBat could not be applied in this case but we used the ARSyN method^51^ instead which can estimate the systematic sources of noise and then correct the data to remove them.

#### RRBS

Initial quality assessment was based on data passing the Illumina Chastity filter. The second quality assessment was based on the reads using the Fastqc quality control tool version 0.10.0. Reads were adaptor- and quality-trimmed using Trim-Galore Software v0.3.4 (http://www.bioinformatics.babraham.ac.uk/projects/trim_galore/) in RRBS paired-end mode, in order to decrease methylation call errors arising from poor quality data. Mapping to the reference genome (GRCm38, mm10) was performed using Bismark v0.10.1^52^ and Bowtie2^50^. The quality of the mapping was inspected using HTSEQ-qa^44^. SAM files were used as input in Bismark to obtain methylation calls. Paired-end mode with no overlap mode was specified. The first four bases from each read were avoided to eliminate M-bias, i.e. deviation from the horizontal line in the mean CpG methylation level for each read position. BedGraph and *.cov files were further considered and analyzed with the BiSeq package^53^. Coverage was inspected before proceeding to smooth the methylation levels (between 0 and 1) per CpG site. Briefly, we firstly defined “frequently covered CpG sites” as those sites that are covered in at least 2/3 of the samples. The frequently covered CpG sites were considered only to define the cluster boundaries and we defined CpG clusters using a maximum distance of 100 bp and at least 20 CpGs. This selection resulted in 1,116,417 CpG sites within CpG clusters, with no threshold on coverage. The extra coverage of unusually high covered sites (95% quantile of the coverage) was eliminated to remove potential biases during the smoothing step introduced by CpGs with exceptional high coverage. Then, the methylation levels were smoothed with a bandwidth of 80 bp as described^53^. Clustering analysis was performed with the methylation estimates of the 20% most variable positions (based on CV), with multidimensional scaling or hierarchical clustering. M-values were obtained after thresholding methylation levels in the interval [0.01, 0.99] to avoid infinite values, as M = log2(b/(1-b)), where b is the constrained methylation level. The final dataset contained a total of 1,116,417 Methylation features.

#### Single-cell RNA-seq

A total of 560 single-cell RNA-seq libraries were mapped with Tophat^54^ to the mouse Ensembl gene annotations and mm10 reference genome. Single-cell libraries with a mapping rate less than 50% and less than 450,000 mapped reads were excluded from any downstream analysis, resulting in 324 single-cells for all subsequent analysis. Cufflinks^55^ version 2.2.1 was used to quantify expression from single-cell libraries using Cuffquant. Gene expression measurements for each single-cell library were merged and normalized into a single data matrix using Cuffnorm. Genes with zero counts in more than 80% of the samples were removed resulting in a data matrix with 9,075 genes.

#### ATAC-seq

Single-cell libraries were mapped with Bowtie^50^ to the mm10 reference genome using the following parameters (bowtie -S -p 2 --trim3 10 -X 2000). Duplicate fragments were removed using Picard (http://picard.sourceforge.net). We considered single-cell libraries that recovered > 5k fragments after mapping and duplication removal. Bulk ATAC-seq replicates were mapped to the mm10 reference genome using the following parameters (bowtie2 -S -p 10 --trim3 10 -X 2000). Peak calling was performed on bulk replicates using HOMER with the following parameters (findPeaks <tags> -o <output> -localSize 50000 - size 150 -minDist 50 –fragLength 0). The intersection of peaks in three biological replicates was performed. A consolidated list of 25,466 peaks was generated from the union of peaks from 0 and 24 hour time-points.

#### Proteomics

Data were searched against a protein sequence database containing all confirmed mouse protein sequences from the Uniprot database (swissprot), common contaminants and reversed using the Andromeda algorithm within the MaxQuant software suite version 1.5.0.0. Mass deviation settings for peptide detection were 20 ppm for the first search and 7 ppm for the main search. IT MS/MS data were searched with a mass accuracy of 0.6 Da. N-terminal acetylation and methionine oxidation were set as variable modifications, carbamidomethylation of cysteine as fixed modification. Unidentified signals, present at a similar retention environment, were matched if at least one run had a positive identification of the peptide sequence by enabling the match between run option within an alignment time window of 30 min. Obtained search results were filtered for reversed database hits on the peptide spectrum match level (1%) and the protein level (2%) and all protein groups with at least 1 razor or unique peptide were initially accepted. For quantitation of proteins over the different timepoints, light protein intensities were extracted from the data file for each protein.

Distant measuring intervals and long gradient times lead to a substantial variation between peak localization and areas of individual LC-MS/MS runs resulting in a huge number of missing values. This issue was addressed by:

a. Alternative LC MS/MS alignment routines. Upon the observation that missing values were not randomly distributed, but associated to particular samples, we believed that a misalignment of chromatograms played a role. We improved the alignment between samples to rescue some of the missing values.
b. RNA-seq data was used as database source protein identification. The rationale is that the mRNAs of expressed proteins should be found within the RNA-seq detected genes. Reducing the size of the protein database to proteins detected by RNA-seq will reduce the number of false hits and lead therefore to more specific data on the pre-B cell proteome. Since proteomic data exhibit a lower coverage of the proteins abundant in a cell, a substantial data loss is not expected.
c. A conservative missing value imputation strategy was applied to log2-transformed data corresponding to 2,527 proteins. Briefly, this strategy discarded proteins with a large number of missing values, considered as not expressed those proteins that were missing either in the whole Control or Ikaros condition, and imputed values in conditions with only 1 out of 3 missings. Samples were normalized by the mean of medians per experimental condition. Proteins with missing values in all the 3 replicates per condition in at least 11 of the 12 conditions were discarded, resulting in 2,396 proteins used for the imputation. Proteins with all missing values in the Control condition were imputed from a Gaussian distribution with mean 50% of the minimum sample value and with standard deviation equal to the median of all within-group standard deviations for all the proteins in the original data. The same procedure was followed for Ikaros condition. Finally, when for a given condition only one of the three biological replicates was missing, the missing value was computed as before but using the mean of the two measured values. The resulting imputed data with no missing values were normalized by TMM. Both imputed and non-imputed datasets are available at the STATegra figshare repository.

#### Metabolomics

For the GC platform, a 13C labeled yeast extract was added as internal standard. A Quality Control (QC) sample was measured every 6 samples. For each of the compounds measured on the GC platform, the labeled compound peak that led to the smallest standard error in the QC samples for that compound, was selected as internal standard. Because the amount of sample was almost completely used for the two analytical platforms no replicate analyses were possible. This meant that the internal standard selection could not be validated using replicate samples as is common practice. For the targeted LCMS method, the optimal internal standards for each metabolite were chosen during optimization and validation of the method. The limited within batch drift effects were corrected using the batch correction approach developed by van der Kloet *et al*^56^.

Four biological batches (batches 9 through 12) were provided to the metabolomics platforms, which were (physically) different from the batches used for mRNA-seq, miRNA-seq and proteomics. Visual inspection showed that samples of batch 11 and 12 were not completely dry. Analysis of some key metabolites and PCA showed batch 12 levels to be outside the general trend in batches 9, 10 and 11 (not shown). Therefore, it was decided to exclude batch 12 from further analysis.

Both analytical platforms show some overlap in the metabolites that were measured. On GCMS 22 metabolites were uniquely quantified and 18 metabolites were quantified uniquely on LCMS, while 18 metabolites were quantified both on GCMS and LCMS, making a total of 58 metabolites. Although the intensity levels of the GCMS and LCMS were rather different a high correlation between the two platforms for most overlapping metabolites was observed. Metabolite levels were log scaled and levels were mean-centered over the three batches 9, 10 and 11. For the metabolites that were measured both on GCMS and LCMS, the LCMS values were selected as this platform is targeted for these types of metabolites.

## Data Records

### Raw data

STATegra multi-omics data have been deposited in different public repositories dedicated to different data types. Table 1 shows a list of the current hosting of raw data files. Moreover, pre-processed data arranged as a data-matrix per omics data-type have also made available at Lifebit (https://lifebit.ai/) site. Additionally, preprocessing scripts and intermediate files are available at the STATegraData GitHub repository.

**Table 1.** Public repositories hosting STATegra multi-omics data.

| Data set | Database and accession |
| --- | --- |
| mRNA-seq | GEO, GSE75417 |
| miRNA-seq | GEO, GSE75394 |
| RRBS | GEO, GSE75393 |
| DNase-seq | GEO, GSE75390 |
| ATAC-seq | GEO, GSE89362 |
| scRNA-seq | GEO, GSE89280 |
| scATAC-seq | GEO, GSE89362 |
| ChIP-seq | GEO, GSE38200 |
| Proteomics | ProteomeXchange,<br>PXD003263 |
| Metabolomics | MetaboLights, MTBLS283 |

### STATegra Knowledge Base

In order to evaluate how to best integrate and semantically map specific prior knowledge and relevant information derived from multiple sources together with heterogeneous experimental data, a STATegra Knowledge network for B-cell differentiation (KB) was developed (applying the BioXM™ knowledge management environment^57^ (https://ssl.biomax.de/stategrakb/). Prior knowledge includes among others relevant molecular elements (genes, proteins, metabolites, etc.), functional information (GO, OMIM, etc.), functional interactions (e.g. protein-protein interaction, transcriptional regulation (e.g. mouse TF-regulatory network), miRNA network, etc.) and information about gene homologs (mouse, rat, human). Also, genome features with coordinates for peak-to-gene associations of NGS data (e.g. mouse genome assembly mm10), metabolic and signal transduction pathways, cell types related to B-cell differentiation as well as ontologies such as the mouse anatomy ontology (MGI) were incorporated. This integrated and dynamically organized knowledge serves as information rich, structured background network. Semantic mapping of experimental data to this background network of prior knowledge enables complex integration and analysis approaches^58^. The STATegra Knowledge Base visualizes multiple omics data types and summarizes information from different layers on top of a network graph. The overlay of experimental data on top of such networks helps interpretation of results as well as validate database predictions.

## Technical validation

### Validation of time course replicability

As a quality control of batch replicability, real time RT-PCR was used to check the impact of Ikaros in gene expression upon induction and reproducibility across time course experiments. RNA from all samples was extracted using RNAbee (AMS Biotechnology (Europe) Ltd) and treated with Turbo DNAse (Life technologies). Bioanalyser technology was used to check the RNA integrity and samples were quantified using a Nanodrop. Changes in the expression of few previously identified Ikaros-responsive genes were analyzed (Figure 5). As expected^34^, early down-regulation of *Igll1* and *Myc*, late down-regulation of *Slc7a5*, *Hk2* and *Ldha*, and up-regulation of *Foxo1* and *Lig4* were consistently observed in the three independently collected time course replicates. Either frozen pellets or RNA samples from the time course experiments and 0h time point collections were sent to the different experimental labs to perform the library preparation for the sequencing.

**Figure 5.**
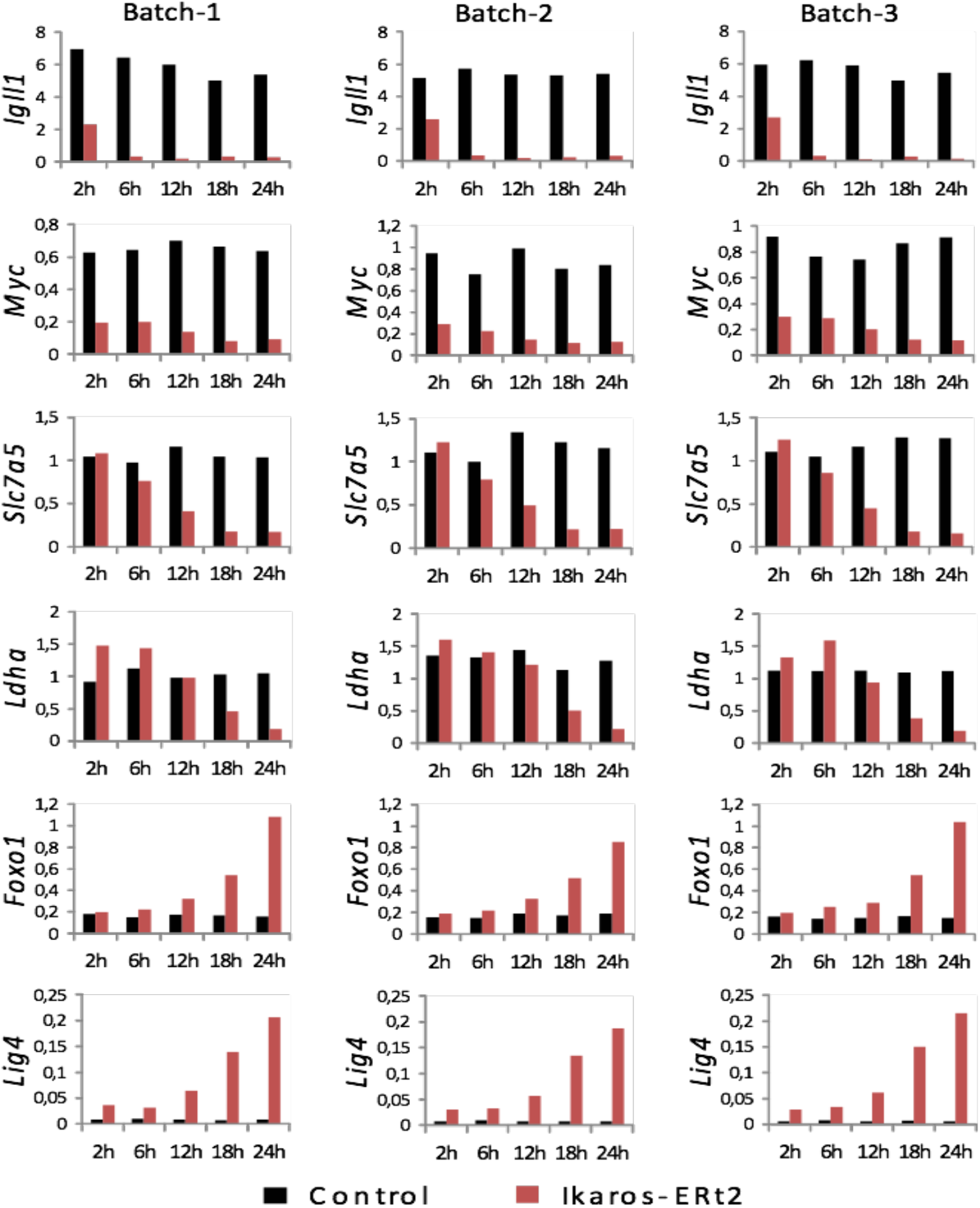
Biomarkers of B3 cell differentiation across three experimental batches.

### Validation of dataset replicability and co-variation structure

To assess the quality of our data we analyzed the correlation values between replicates of the same condition and compared to correlation values when samples belonging to different conditions were compared (Figure 6A). We applied this analysis to RNA-seq, miRNA-seq, Dnase-seq, Methyl-seq, proteomics and metabolomics. ChIP-seq and ATAC-seq data were excluded as only two replicates were available in each case. Also, single-cell data was excluded from this analysis, as the zero-inflated nature of the technologies makes correlation analysis meaningless. We found that, for all technologies, biological replicates had very high correlation (> 0.9, Figure 6A), in general higher than the correlations among samples of different experimental conditions that also displayed a wider range of values. This result reflects the time course nature of the experimental design, where closer time points have higher correlations than distant time points.

**Figure 6.**
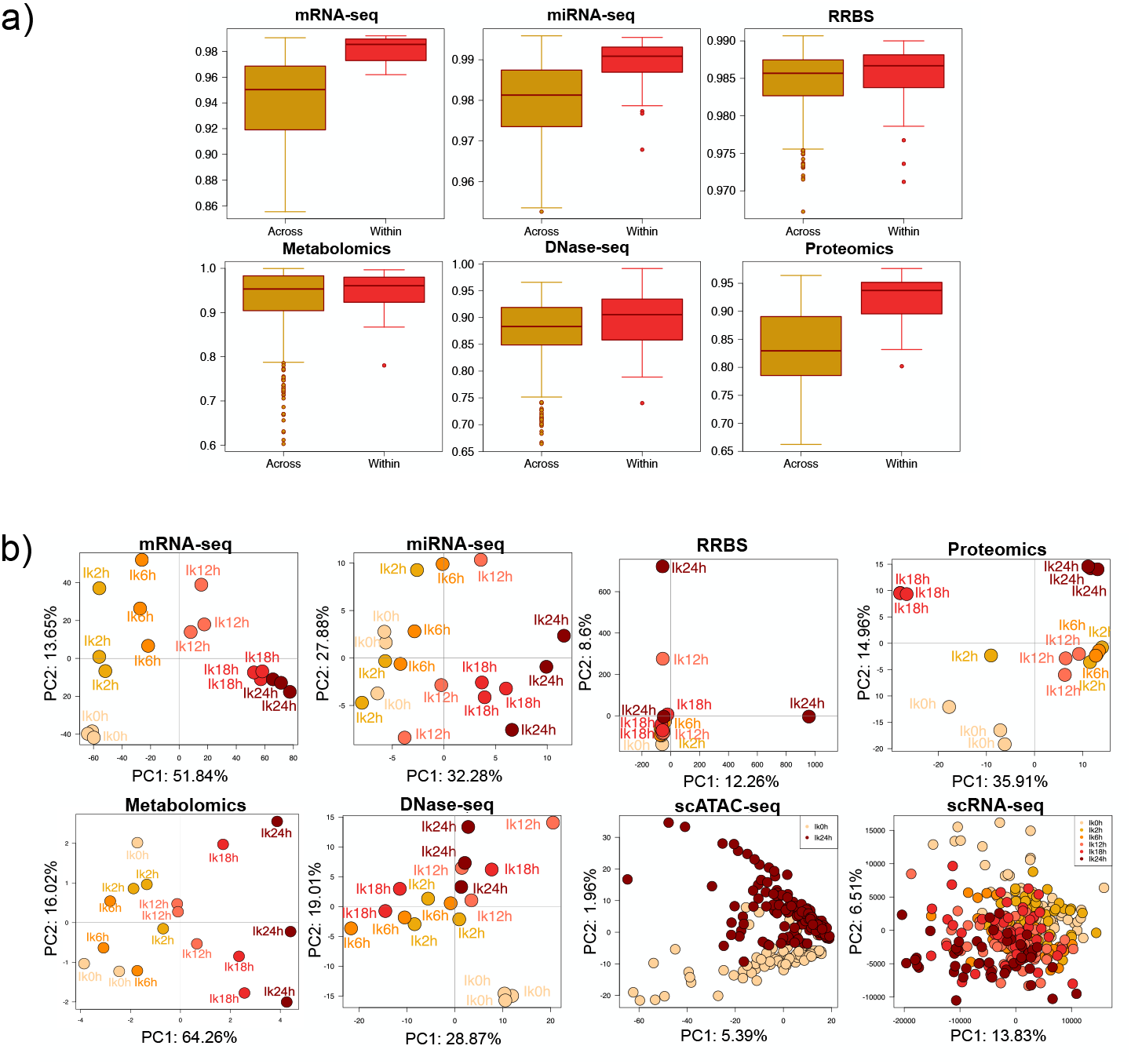
Quality control of STATegra multi-omics data. A) Distribution of pair-wise correlation values for samples belonging to different (Across) or the same (Within) experimental conditions. B) PCA analysis. Only the Ikaros series is shown. Data were preprocessd as described in Methods. Time progression is represented by an increasingly darker red color.

To further validate data and to understand whether the different omics measurements captured the dynamics of B-cell differentiation and/or had a similar co-variation structure, we ran Principal Component Analysis for all datasets (Figure 6B). In general, the different multiomics datasets show PCA plots that recapitulate the time progression of our inducible system. A well spread temporal progression on the first PC was observed for mRNA-seq, miRNA-seq and scRNA-seq data, being RNA-seq the dataset with the most consistent progression signal. Metabolomics and Proteomics showed a two-stage pattern, with samples from 0h-12h hours clustering at negative values, and samples at 18h-24h clustering at positive values of the first and second PC, respectively. DNase-seq showed a noisier, but distinguishable distribution of the temporal signal at PC2, while RRBS is the only dataset with an unclear temporal pattern. For scATAC-seq, only two time points were measured and cells nicely separated on the second PC. This analysis reveals that multi-omics datasets consistently described the progression of B-cell differentiation but also that the different omics technologies present different noise levels. Interestingly, dynamic patterns are slightly different for nucleic acids and proteins and metabolites, possibly indicating a later response of these with respect to the transcriptional change.

### Multi-layer data example

In order to illustrate the consistency of the STATegra multi-layer data, we analyzed values for the *lactate dehydrogenase A* gene (Figure 7). LDHA catalyzes pyruvate to lactate conversion in the final step of anaerobic glycolysis (Figure 7A). *Ldha* is one of known Ikaros target genes^34^ and downregulated upon Ikaros-induced differentiation of the B3 cell line (Figure 5). The STATegra footprint data confirmed that Ikaros binds to the promoter region of the *Ldha* gene (Figure 7B) while the promoter DHS signal, mRNA and protein levels were downregulated as cells progressed towards the pre-BII stage (Figure 7C). We obtained confirmed microRNAs targeting the *Ldha* transcript 3’UTR from the mirWalk database^59^ and identified four microRNAs with a strong negative correlation with *Ldha* expression levels (Figure 7C). One of these microRNAs, mir449a-5p, has been reported to bind and regulated *Ldha* in human cells^60^. Finally, STATegra data indicated a general downregulation of glycolysis (Figure 7D), as previously described, and confirmed a decrease in pyruvate and lactate at mid-late time points of differentiation (Figure 7A). In summary, the STATegra data recapitulates known metabolic-switch observations in the B3 system and showed a consistent pattern of change across regulatory layers.

**Figure 7.**
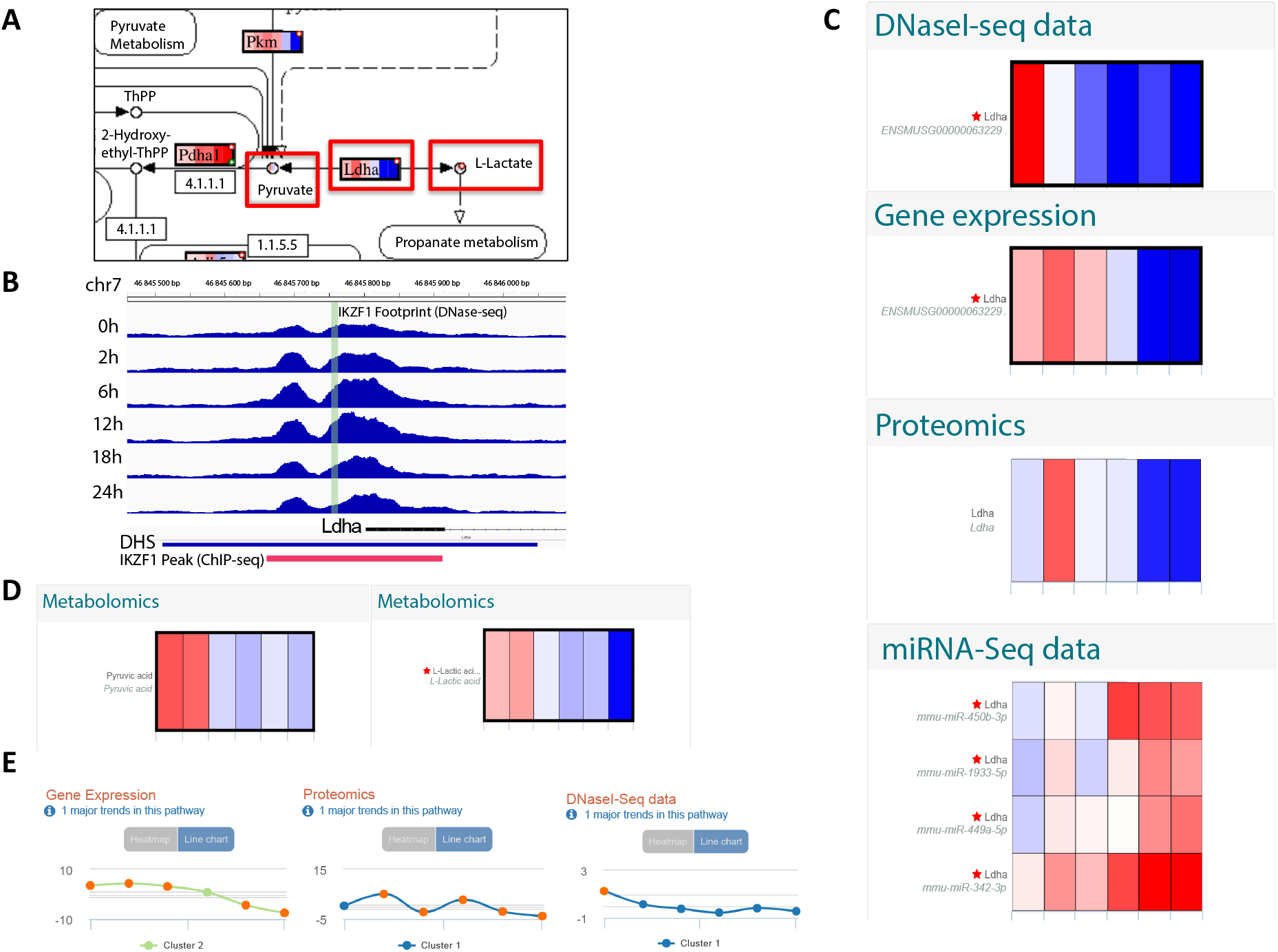
STATegra data for lactate dehydrogenase A. A) LDHA reaction at glycolysis. B) Promoter regions of the *Ldha* gene showing a DSH and IKZF1 footprint identified by DNase-seq. Only values for the Ikaros-induced time course are shown. In red, the IKZF1 ChIP-seq peak region. C-E: Paintomics^27^ representation of Ldha data. Boxes display mean log2FC values between Ikaros and Control at 0,2,6,12,18 and 24 hours after Ikaros induction. Red indicates up-regulation, blue indicates down-regulation. C) Ldha data for DNase-seq, RNA-seq, Proteomics, and miRNA-seq. miRNA-Ldha target data was predicted by at least 5 algorithms in the mirWalk^59^ database. D) STATegra log2FC values for pyruvate (left) and lactate (right). E) Major Gene Expression, Proteomics, and DNase-seq trends for glycolysis pathway computed by Paintomics^27^.

## Acknowledgements

This work has been funded by the European Union Seventh Framework Programme [FP7/2007–2013] under the grant agreement 306000-STATegra.

The authors declare no competing interests

## Author contributions

DGC: Conceived the study, contributed to RNA-seq data generation, performed preprocessing and validation of multiple omics datasets

ST: Performed preprocessing of multiple omics datasets, created the initial draft of the paper

IFV: Generated B3 cell line cultures for multi-omics profiling

RNR: Performed DNase-seq, ATAC-seq, scATA-seq and scRNA-seq experiments. Contributed to data preprocessing and drafting of the manuscript.

CC: Performed RRBS and microRNA-seq experiments

AS: Performed Proteomics experiments and contributed to data preprocessing

TR: Performed Metabolomic experiments and contributed to data preprocessing

VvSP: Developed STATegra KnowledgeBase portal

FM: Performed preprocessing of RRBS data

JRU: Contributed to the preprocessing of miRNA-seq and RRBS data AGG: Contributed to the preprocessing of miRNA-seq and RRBS data

TC: Contributed to generating B3 cell line cultures for multi-omics profiling

LC: Contributed to generating B3 cell line cultures for multi-omics profiling

ZL: Contributed to generating B3 cell line cultures for multi-omics profiling

GD: Contributed to generating B3 cell line cultures for multi-omics profiling

FvdK: Contributed to Metabolomics preprocessing

ACH: Obtained Metabolomics data and performed initial measurements

LBN: Contributed to the analysis of multiomics dataset

VL: Contributed to data analysis

IT: Contributed to the supervision of analyses

ML: Contributed to the supervision of analyses

DM: Supervised STATegra KnowledgeBase development

JAW: Supervised data preprocessing

TH: Supervised Metabolomics data generation

AI: Supervised Proteomics data generation

EB: Supervised miRNA-seq and RRBS data generation

AM: Supervised DNase-seq and single-cell data generation

MM: Supervised cell culture experiments and validation

JT: Conceived the study and supervised analyses

AC: Conceived and coordinated the study supervised analyses and drafted the manuscript.

## Notes

http://www.stategra.eu/

https://ssl.biomax.de/stategrakb/

